# A novel antimicrobial alternative of microbiota metabolic product deoxycholic acid controls chicken necrotic enteritis

**DOI:** 10.1101/215640

**Authors:** Hong Wang, Juan D. Latorre, Mohit Bansal, Bilal Al-Rubaye, Guillermo Tellez, Billy Hargis, Xiaolun Sun

**Author notes:** To whom correspondence should be addressed: Center of Excellence for Poultry Science, 1260 W Maple St. O-409, Fayetteville, AR 72701.

## Abstract

Necrotic enteritis (NE) caused by *Clostridium perfringens* infection has reemerged as a prevalent poultry disease worldwide due to reduced usage of prophylactic antibiotics. The lack of alternative antimicrobial strategies to control this disease is mainly due to limited insight into NE pathogenesis, microbiome relationships, and host responses. Here we reported that the metabolic byproduct of microbial metabolism of bile acids to deoxycholic acid (DCA), at as low as 50 μM, inhibited 82.8% of *C. perfringens* growth in Tryptic Soy Broth (P < 0.05). Sequential *Eimeria maxima* and *C. perfringens* challenge strongly induced NE, severe intestinal inflammation, and body weight (BW) loss in broiler chickens. These negative effects were diminished by 1.5 g/kg DCA diet. At the cellular level, DCA alleviated NE-associated ileal epithelial death and lamina propria immune cell apoptosis. Interestingly, DCA reduced *C. perfringens* invasion into villi without significantly altering the bacterial luminal colonization. Molecular analysis showed that DCA reduced inflammatory mediators of *Infγ, Litaf (Tnfα), Il1β,* and *Mmp9* mRNA accumulation in ileal tissue. Mechanically, *C. perfringens* induced elevated expression of inflammatory cytokines of *Infγ, Litaf,* and *Ptgs2* (COX-2 gene) in chicken splenocytes. Inhibiting the COX signaling by aspirin attenuated INFγ- or TNFa-induced inflammatory response in the splenocytes. Consistently, chickens fed 0.12 g/kg aspirin diet resisted against NE-induced BW loss, ileal inflammation, and villus apoptosis. In conclusion, microbial metabolic product DCA prevents NE-induced BW loss and ileal inflammation through curbing inflammatory response. These novel findings could serve as a stepping-stone for developing next generation antimicrobial alternatives against NE.

## Significance Statement

Widespread antimicrobial resistance has become a serious challenge to both agricultural and healthcare industries. Withdrawing antimicrobials without effective alternatives exacerbates poultry productivity loss at billions of dollars every year, caused by intestinal diseases, such as coccidiosis- and *C. perfringens-induced* NE. This study used the endogenous microbial metabolic secondary bile acid product DCA to control NE and to improve chicken growth performance through modulating host response. These findings have opened new avenues for developing next-generation antimicrobial free regiments.

## Introduction

Antimicrobial resistance is one of the emerging challenges requiring immediate and sustainable counter-actions from agriculture to healthcare (1). Increasing antimicrobial resistance has caused the emergence of multiple drug-resistant microbes or “superbugs”. Recently, a “superbug” of an *Escherichia coli* strain resistant to the last resort antibiotic, Colistin, was reported in USA (2). Overuse of antimicrobial agents in medical and agricultural practice is contributing to exacerbating the episodes of emerging antimicrobial resistant microbes (1). Withdrawing antimicrobials in poultry production, however, has caused new problems for the poultry industry by reducing production efficiency and increasing diseases, such as *Eimeria maxima*- and *Clostridium perfringens-induced* necrotic enteritis (NE) (3). Clinical signs of NE include watery to bloody (dark) diarrhea, severe depression, decreased appetite, closed eyes, and ruffled feathers. Dissection of dead or severely ill birds shows that the intestine is often distended with gas, very friable, and contains a foul-smelling brown fluid, with clearly visible necrotic lesions (4). Although progress has been made toward understanding risk factors influencing the outcome of NE such as *C. perfringens* virulence, coccidiosis, and feed (5), few effective non-antimicrobial strategies are available.

The human and animal intestine harbors up to trillions of microbes and this intestinal microbiota regulates various host functions such as the intestinal barrier, nutrition and immune homeostasis (6-8). The enteric microbiota regulates granulocytosis and neonatal response to *Escherichia coli K1* and *Klebsiella pneumoniae* sepsis (9), suggesting the key role of the microbiota in protecting the host against systemic infection. At the gut level, fecal transplantation was reported decades ago to prevent *Salmonella infantis* chicken infection (10). More recently, microbiota transplantation has shown tremendous success against recurrent *Clostridium difficile* infection (11) and *Clostridium scindens* metabolizing secondary bile acids have been shown to inhibits *C. difficile* infection (12). Bile acids synthesized in the liver are released in the intestine and metabolized by gut microbiota into final forms of secondary bile acids (13). Bile acids, particularly the secondary bile acid DCA, are associated with a variety of chronic diseases, such as obesity, diabetes, and colorectal tumorigenesis (14, 15). Recently, we found that mouse anaerobes and their metabolic product DCA prevented and treated *Campylobacter jejuni*-induced colitis in germ-free mice through attenuating host inflammatory signaling pathways (16).

Cyclooxygenases (COX)-catalyzed prostanoids regulate various activities including cell proliferation, apoptosis and migration (17), gastrointestinal secretion (18), body temperature (19), inflammation (20), and pain sensation (21). COX-1 and COX-3 (gene *Ptgs1*, alternative splicing) constitutively expressed are important for intestinal integrity. Inducible COX-2 (*Ptgs2*) activity is associated with various inflammatory diseases including inflammatory bowel disease (22) and radiation-induced small bowel injury (23). COX-2 increases gut barrier permeability and bacterial translocation across the intestinal barrier (24, 25). Paradoxically, COX-2 enhances inflammation resolution through prostaglandin D2 (26). Although non-selective COX inhibitor aspirin is used to prevent various chronic diseases, it inflicts intestinal inflammation to the healthy intestine (27).

Currently, limited knowledge is available on the relationship between NE pathogenesis, the microbiome, and host inflammatory response. Here, we hypothesize that the microbiota metabolic product DCA attenuates NE. Our findings demonstrated that DCA decreases NE-induced BW loss, intestinal inflammation, *C. perfringens* invasion, and villus death. Blocking the inflammatory downstream target COX signaling pathways by aspirin reduces NE-induced intestinal inflammation. These findings could serve as a stepping-stone for the development of new antimicrobial free prevention and therapeutic strategies against NE.

## Results

### DCA prevents *C. perfringens in vitro* growth

We previously reported that the secondary bile acid DCA prevents and treats *C. jejuni-induced* intestinal inflammation in germ-free mice (28). This secondary bile acid also inhibits *C. difficile in vitro* growth (12). Since *C. difficile* and *C. perfringens* are in the same genus, we reasoned that DCA would prevent *C. perfringens* growth. To test this hypothesis, we implemented *in vitro* inhibition experiments, in which *C. perfringens* was inoculated in Tryptic Soy Broth (TSB) with sodium thioglycollate under anaerobic condition. The TSB was also added with various concentrations of bile acids, including conjugated primary bile acid taurocholic acid (TCA), primary bile acid cholic acid (CA), and secondary bile acid DCA. Notably, DCA inhibited *C. perfringens* growth at 0.01 (−33.8%) and 0.05 mM (−82.8%, clear broth), respectively, compared to control, while TCA (−16.4%) and CA (−8.2%) barely prevented the bacterial growth (cloudy broth) even at 0.2 mM (Figure 1A and B). We then examined if other secondary bile acids were also bacteriostatic in TSB. Interestingly, *C. perfringens* growth was only mildly inhibited by lithocholic acid (LCA; −22.6 and −23.8%) and ursodeoxycholic acid (UDCA; −10.0 and −25.3%) at 0.2 and 1 mM, respectively (Figure 1C and D). These results suggest that the secondary bile acid DCA effectively curbs *C. perfringens in vitro* growth.

**Figure 1.**
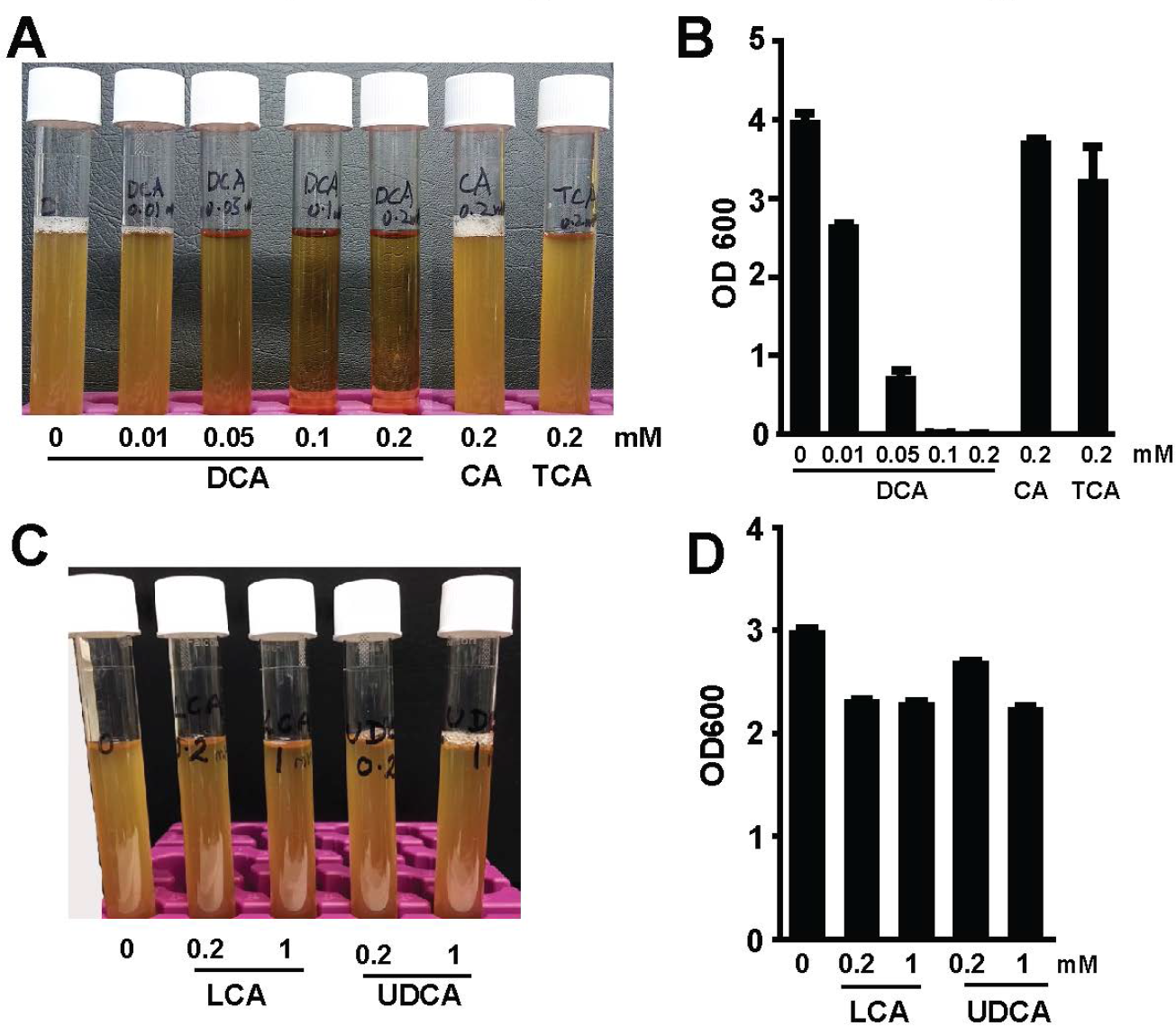
DCA inhibits *C. perfringens in vitro* growth. *C. perfringens* (10^3^ CFU) were inoculated into TSB supplemented with various concentrations of conjugated primary bile acid TCA, primary bile acid CA, and secondary bile acid DCA. (A) Image of bile acids did (clear broth) or did not (cloudy broth) inhibited *C. perfringens* growth. (B) OD600 reading of the broth in A. (C) Image of bile acids did not inhibit *C. perfringens* growth. (B) OD600 reading of the broth in C. All graphs depict mean ± SEM. Results are representative of 3 independent experiments.

### DCA prevented NE-induced productivity loss

To further address whether DCA reduces coccidia *E. maxima*- and *C. perfringens*-induced necrotic enteritis (NE) in birds, we fed day-old broiler chicks with 1.5 g/kg CA or DCA diets and growth performance of body weight (BW) gain were measured. To induce NE, the birds were infected with 20,000 sporulated oocysts/bird *E. maxima* at 18 days of age and then infected with 10^9^ CFU/bird *C. perfringens* at 23 and 24 days of age. Notably, DCA (solid black bar) but not CA (straight line bar) diet promoted bird daily BW gain during 0-18 days of age compared to birds fed control diets (open bar, Figure 2 A). Body weight gain was impaired in birds infected with *E. maxima* (Em) at 18-23 days of age (coccidiosis phase). Subsequent *C. perfringens* infection further drove NE control birds (dotted bar) into BW loss at 23-26 days of age (NE phase). Remarkably, DCA prevented productivity loss at coccidiosis and NE phases compared to the NE control birds. Interestingly, the primary bile acid CA diet attenuated body weight loss at NE phase but failed at coccidiosis phase compared to the NE control birds.

**Figure 2.**
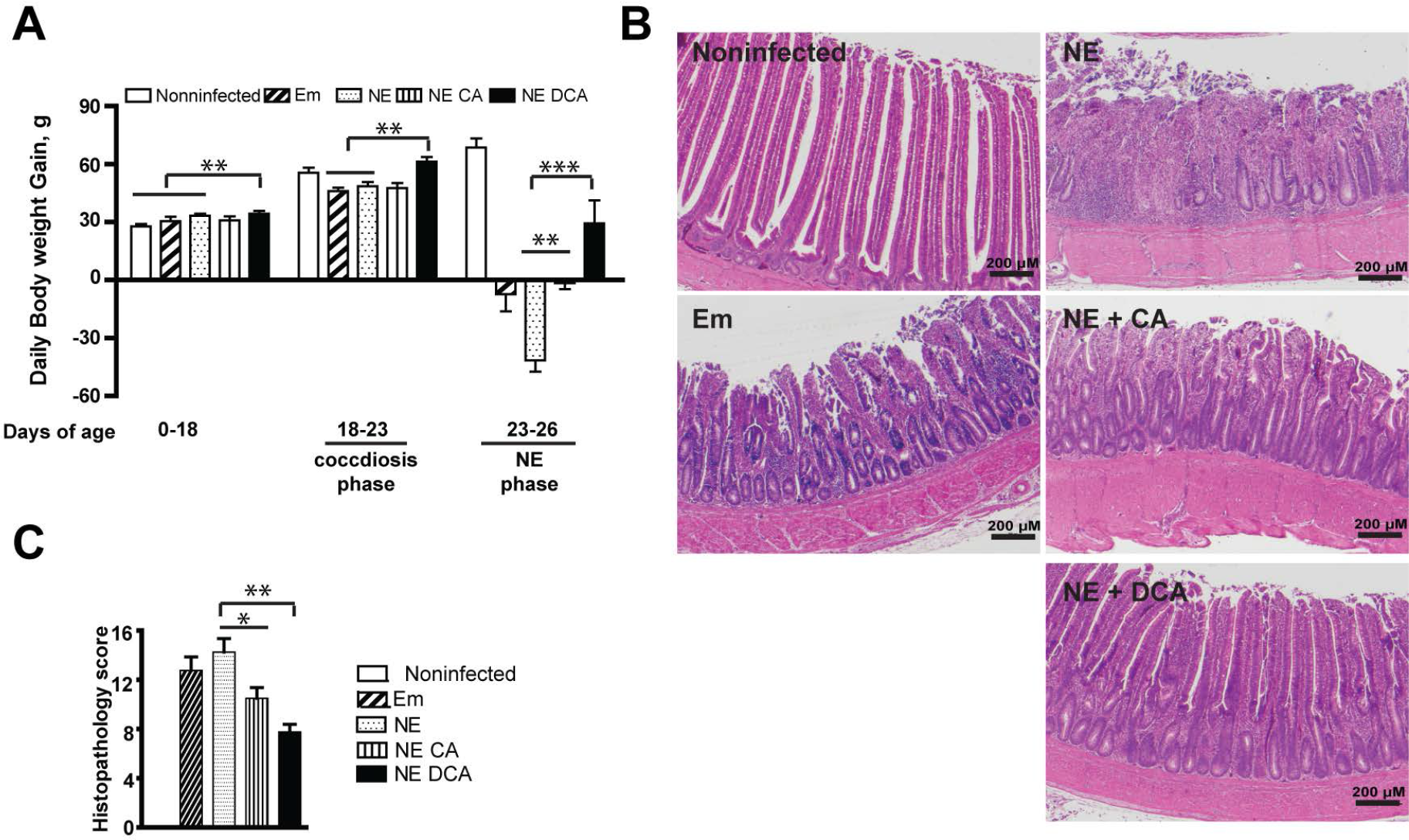
DCA attenuates NE-induced productivity loss and histopathology. Cohorts of 13 broiler chickens were fed basal, 1.5 g/kg CA, or DCA diets. The birds were infected with *E. maxima* and *C. perfringens* at 18 and 23 days of age, respectively. The birds were sacrificed at 26 days of age. (A) Bird growth performance of daily body weight gain. (B) H&E staining showing representative intestinal histology images. (C) Quantification of histological intestinal damage score. Scale bar is 200 μm. All graphs depict mean ± SEM. *, P<0.05; **, P<0.01; ***, P<0.001. Results are representative of 3 independent experiments.

### DCA prevented NE-induced histopathology

Coccidiosis and NE induce severe intestinal inflammation. To have a thorough insight into DCA impact on NE pathogenesis, we collected intestinal tissue at upper ileum as Swiss rolls, processed with H&E staining, and performed histopathology analysis. Notably, *E. maxima* infection induced severe intestinal inflammation (ileitis) as seen by immune cell infiltration into lamina propria, crypt hyperplasia, and mild villus height shortening compared to uninfected birds (Figure 2B). Furthermore, NE control birds suffered worse ileitis as seen by necrosis and fusion of villi and crypt, massive immune cell infiltration, and severe villus shortening. Notably, DCA diet dramatically attenuated NE-induced ileitis and histopathology score (Figure 2B and C), while CA reduced NE-induced ileitis and histopathology score. These results indicate that DCA promotes growth performance and resists against coccidiosis- and NE-induced BW loss and severe ileitis and histopathology.

### DCA attenuates NE-induced intestinal cell necrosis and apoptosis

Healthy intestinal epithelial cells have polarity (29) and their nuclei are located toward the basal membrane (30), while stressed dying (apoptosis or necrosis) cells lose polarity (31) and their nuclei disperse from basal to apical membranes (32). We then sought to examine whether cell death is relevant in DCA-attenuating NE-induced ileitis. Since it is difficult to find reliable chicken antibodies to detect apoptosis or necrosis in chicken histology slides, we first resorted to classical histological analysis under high magnification. Consistently, the epithelial nuclei (dark blue) in heathy control bird villi were distributed close to the basal membrane (at the right side of the yellow dash line, Figure 3A lower panel left). In contrast, the nuclei in inflamed villi epithelial cells of Em and NE birds were scattered from basal to the apical membranes, indicating epithelial cell death in villi of those birds. Notably, the DCA diet prevented epithelial cell nucleus translocation to apical side, suggesting cell death reduction. To further characterize the villus cell death, we used terminal deoxynucleotidyl transferase (TdT) dUTP Nick-End Labeling (TUNNEL) assay, which detects later stage of cell apoptosis. Consistent with histopathology results, coccidiosis and NE induced massive scatter (Em birds) or concentrated (NE birds) apoptosis cells (green dots) in villus lamina propria, however, cellular apoptosis was attenuated in the DCA treatment. These results indicate that DCA resists against coccidiosis- and NE-induced cell death in villus epithelial and lamina propria cells.

**Figure 3.**
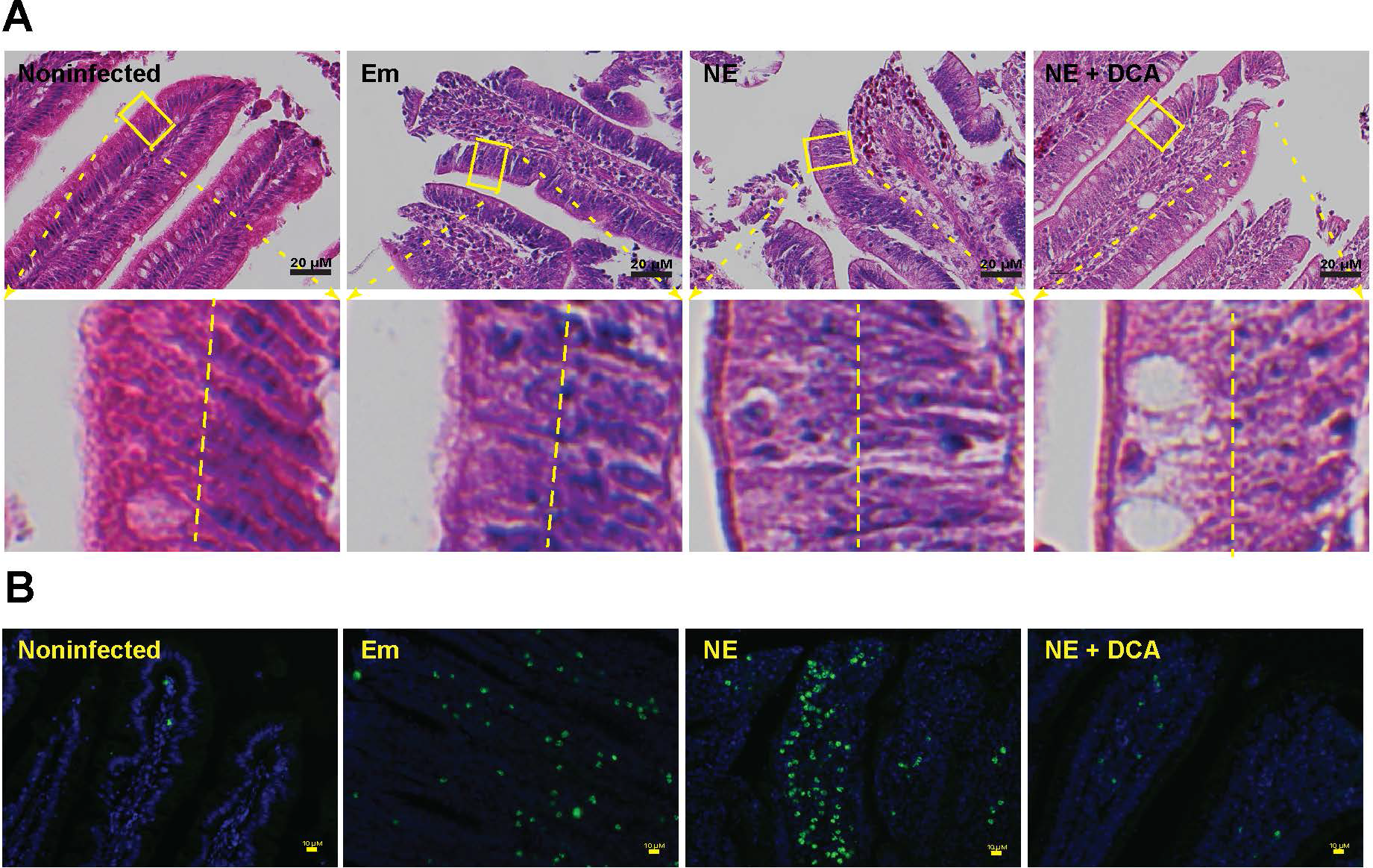
DCA attenuates NE-induced intestinal cell death and apoptosis. Cohorts of 13 broiler chickens were fed different diets and infected as in Figure 2. (A) Representative intestinal cell death (deviated nuclei) using H&E staining. (B) Representative villus cell apoptosis (green) using TUNEL assay. Scale bar is 20 μm (A) and 10 μm (B). Results are representative of 3 independent experiments.

### DCA reduces *C. perfringens* invasion and NE-induced inflammatory response

Given DCA inhibited *C. perfringens* growth *in vitro*, it is logic to reason that DCA might also reduce *C. perfringens* intestinal overgrowth in NE birds. To examine this possibility, we collected ileal digesta extracted total DNA and measured *C. perfringens* colonization level in the intestinal lumen using real-time PCR of *C. perfringens* 16S rDNA. Consistently, coccidiosis and NE were associated with increased luminal *C. perfringens* colonization compared to uninfected birds (Figure 4A). Surprisingly, ileal luminal *C. perfringens* load in DCA birds was not significantly different from NE control birds, while bird productivity and histopathology were strikingly distinct between the two groups of birds (Figures 2A-C). We, therefore, reasoned that the pathogen invasion into tissue was the main driving factors of NE pathogenesis, but not the pathogen luminal colonization level. Using a fluorescence *in situ* hybridization (FISH) technique, we found that while *C. perfringens* was present deeply in the inflamed villus and crypt lamina propria of NE control birds, the bacterium was barely detectable in the ileal tissue of DCA birds (Figure 4B). Because DCA reduced *C. perfringens* invasion and intestinal inflammation, we evaluated the impact of DCA on various proinflammatory mediators in ileal tissue using Real-Time PCR. Notably, *C. perfringens* strongly induced inflammatory *Infγ, Litaf* (*Tnfα*), *Il1β*, and *Mmp9* mRNA accumulation in chicken ileal tissue, an effect attenuated by 51, 82, 63 and 93%, respectively, in DCA fed chickens (Figure 4C).

**Figure 4.**
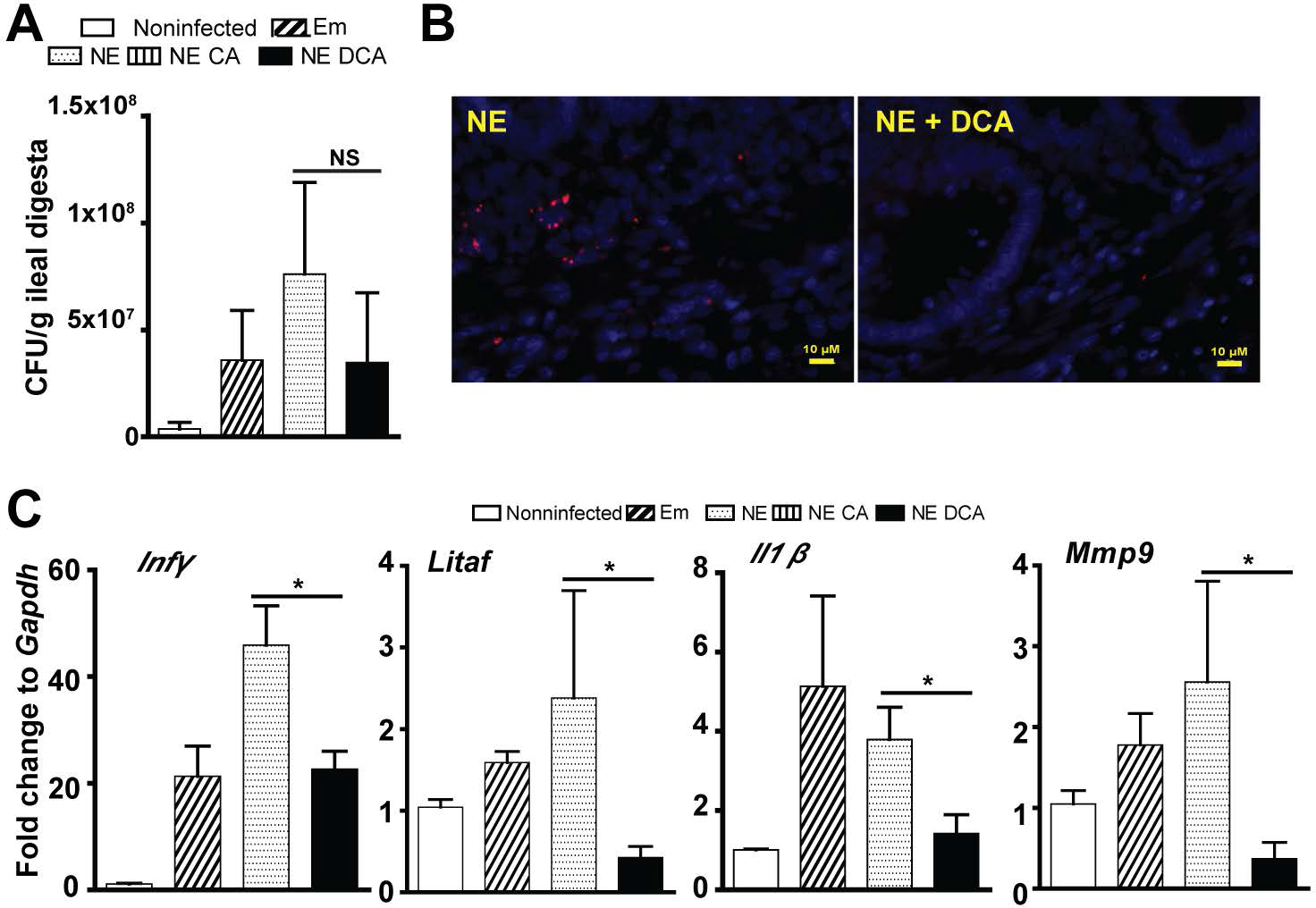
DCA reduces *C. perfringens* invasion and inflammatory response. Cohorts of 13 broiler chickens were fed different diets and infected as in Figure 2. (A) Luminal *C. perfringens* colonization level quantified by 16s RNA real-time PCR. (B) Presence of C. *perfringens* (red dots) in ileal sections of NE birds, detected using fluorescence in situ hybridization (FISH) assay. (C) Ileal *Infγ, Litaf (Tnfα), Il1β*, and *Mmp9* mRNA qPCR fold change relative to uninfected birds and normalized to *Gapdh.* All graphs depict mean ± SEM. NS, not significant; *, P<0.05. Results are representative of 3 independent experiments.

### COX inhibitor aspirin alleviates *C. perfringens-induced* inflammatory response in splenocytes

Inflammatory events shape intestinal diseases and targeting the inflammatory response attenuates disease progress such as inflammatory bowel disease (33) and campylobacteriosis (34). To dissect how the host inflammatory response is involved in NE-induced ileitis, we then used a primary chicken splenocyte cell culture system (35). After isolation from 28-day old chickens, the splenocytes were infected with *C. perfringens* (MOI 100) for 4 hours. Notably, *C. perfringens* increased inflammatory mediators of *Infγ, Litaf, Il1β, Mmp9* and *Ptgs2* (protein COX-2) mRNA accumulation by 1.54, 1.69, 76.47, 1.72 and 8.65 folds, respectively, compared to uninfected splenocytes (Figure 5A). COX-2 are important mediators in the inflammatory response (22). We then used COX inhibitor aspirin towards *C. perfringens-infected* chicken splenocytes, but no inhibition of inflammatory gene expression was observed (data not shown). We then used inflammatory cytokines of recombinant murine INFγ and TNFα to challenge splenocytes in the presence of aspirin. Remarkably, aspirin reduced INFγ-induced inflammatory gene expression of *Infγ, Litaf, Il1β*, and *Mmp9* by 41, 27, 42, and 45%, respectively (Figure 5B). Similarly, aspirin reduced TNFα-induced inflammatory gene expression of *Infγ, Litaf*, and *Mmp9* by 49, 53, and 27%, respectively (Figure 5C). These data indicate that *C. perfringens* induces inflammatory cytokines and COX-2 and inhibiting COX signaling by aspirin reduces the cytokine-induced inflammatory response, suggesting that aspirin poses protection potential against NE detrimental effects.

**Figure 5.**
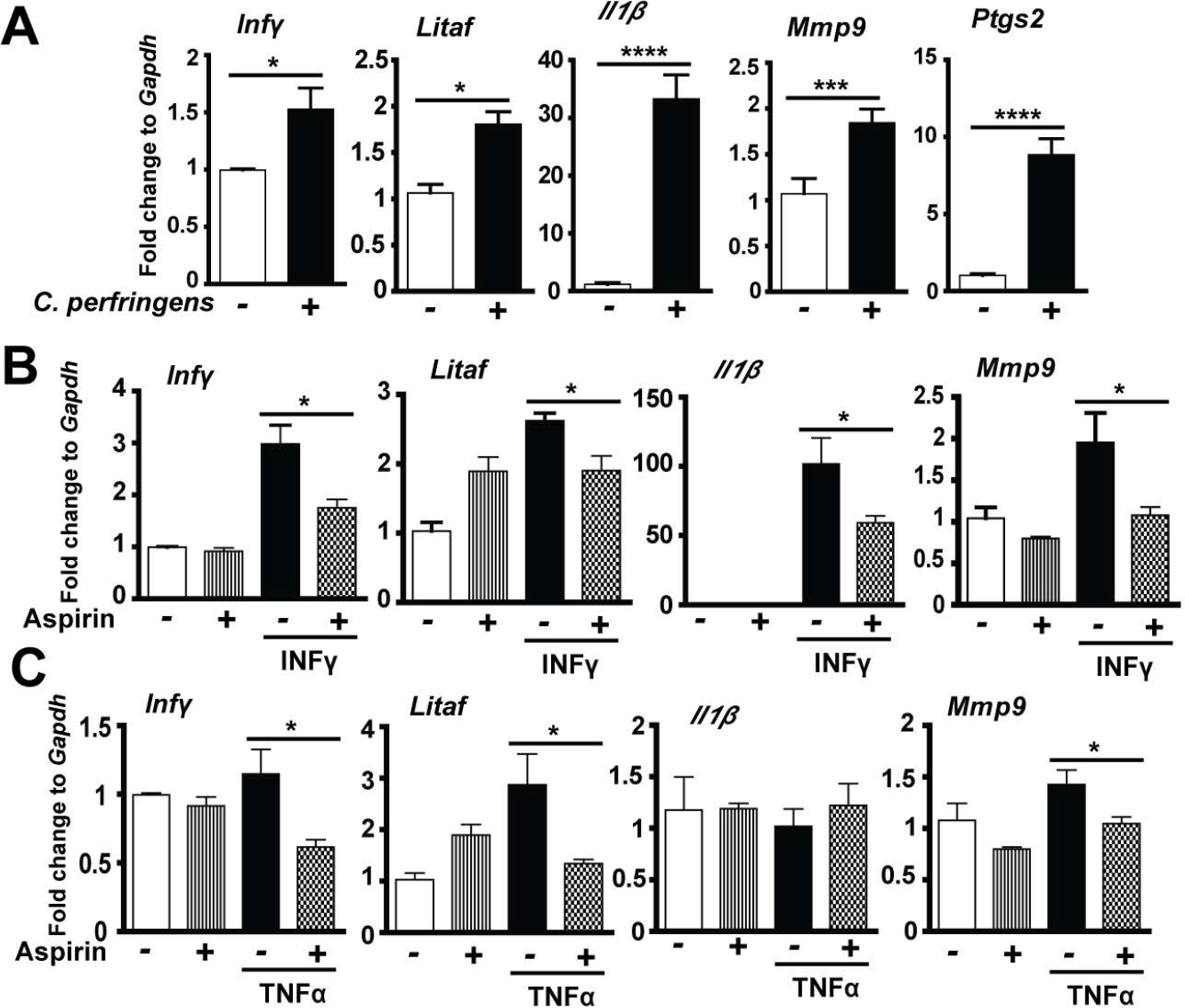
COX inhibitor aspirin alleviates *C. perfringens-induced* inflammatory response in chicken splenocytes. Splenocytes isolated from broiler chickens were infected with *C. perfringens* (MOI 100) for 4 hr or stimulated with murine Infγ (1 μg/ml) or TNFα (5 ng/ml) for 2 hr in the presence of 1.2 mM aspirin. RNA was extracted, reverse-transcribed, and quantified using a Bio-Rad 384 PCR platform. (A) *Infγ, Litaf, Il1β, Mmp9,* and *Ptgs2* mRNA fold change normalized to *Gapdh.* (B) Gene expression fold change in the presence of Infγ and aspirin. (C) Gene expression fold change in the presence of TNFa and aspirin. All graphs depict mean ± SEM. *, P<0.05; ***, P<0.001. Results are representative of 3 independent experiments.

### Aspirin attenuates NE-induced productivity loss, histopathology and apoptosis

To functionally assess the protective effect of aspirin against NE-induced ileitis, broiler chickens fed with 0.12 g/kg aspirin diet (ASP) were infected with *E. maxima* and *C. perfringens* as describe before. Interestingly, ASP birds grew slower compared to control diet birds during 0-18 days of age (Figure 6A). This is because aspirin inhibit all COX isoforms and COX-1 and -3 are important for intestinal homeostats and growth. Notably, ASP attenuated NE-induced BW loss by 60% in NE phase of 23-26 days of age, while no difference between ASP and NE birds in coccidiosis phase of 18-23 days of age. To further dissect the underlying cellular mechanism, we then used a histopathology analysis. Consistently, ASP attenuated NE-induced intestinal inflammation and histopathological score (Figures 6B and C). ASP also reduced NE-induced immune cell apoptosis in villus lamina propria (Figure 6D). These data suggest that aspirin attenuates NE-induced BW loss, intestinal inflammatory response, and villus cell death.

**Figure 6.**
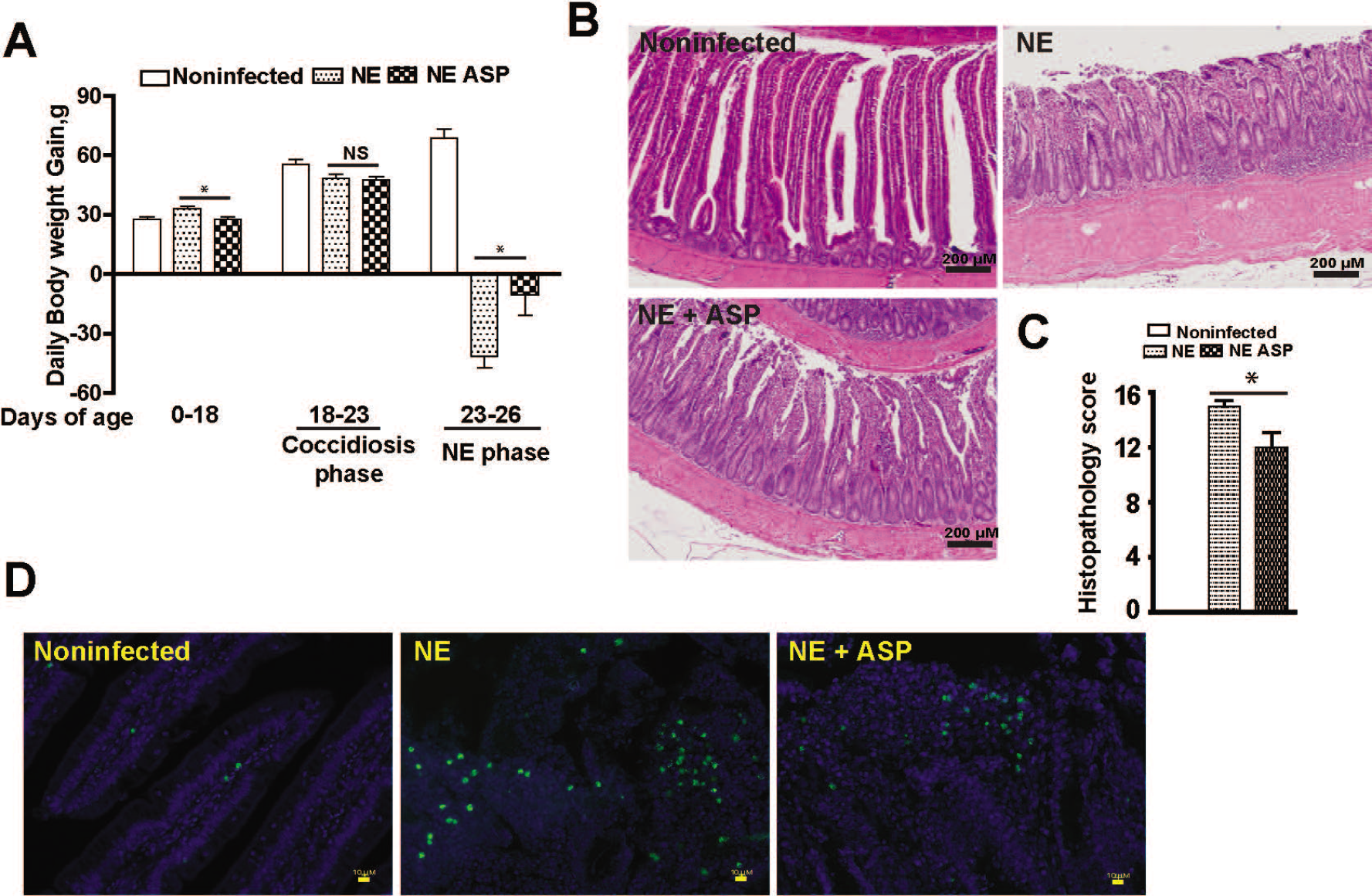
Aspirin attenuates NE-induced productivity loss, histopathology, and apoptosis. Cohorts of 13 broiler chickens were fed basal and 0.12 g/kg aspirin diet. The birds were infected and sampled as in figure 1. (A) Bird growth performance of daily body weight gain. (B) H&E staining showing representative intestinal histology images. (C) Quantification of histological intestinal damage score. (D) Representative cell apoptosis (green) using TUNEL assay. Scale bar is 200 μm in B and 10 μm in C. All graphs depict mean ± SEM. *, P<0.05; **, P<0.01; ***, P<0.001. Results are representative of 2 independent experiments.

## Discussion

Although NE has reemerged as a prevalent poultry disease worldwide in the antimicrobial free era (3), the lack of comprehensive molecular mechanism insight into NE severely hinders the development of antimicrobial alternatives to control this disease (36). Many virulent factors of *Eimeria* and *C. perfringens* are identified but few findings are effective to control NE in poultry production (5, 37), suggesting that we have overlooked important players/factors in NE pathogenesis, such as microbiome and host response. Here we reported that microbiota metabolic product DCA dramatically attenuates chicken NE by reducing chicken inflammatory response. These new findings pave the path for exploring novel antimicrobial alternatives to control NE.

It is a relatively new concept to manipulate microbiota and its metabolic products against infectious diseases. Fecal transplantation was used in chickens decades ago to prevent S. *infantis* infection (10). Microbiome plays an important role in susceptibility to *C. difficile* infection (38). Anaerobe *C.* scindens-transformed secondary bile acids prevent *C. difficile* germination and growth (12). Lithocholic acid and DCA but not primary bile acid CA inhibit *C. difficile* vegetable growth and toxin production (12, 39). However, whether secondary bile acids prevent or treat *C. difficile* infection in human or animal models is still unknown. We recently found that orally gavaging DCA attenuates *C. jejuni*-induced intestinal inflammation in germ-free mice (28). Based on the knowledge, we reasoned that DCA may prevent *E. maxima*- and *C. perfringens-induced* chicken NE. Indeed, dietary DCA but not CA prevents NE and its associated productivity loss. The reduction of ileitis is coupled with reduced *C. perfringens* invasion and intestinal inflammation and cell death. Intriguingly, DCA failed to significantly reduce *C. perfringens* ileal colonization, suggesting that the mechanism of DCA action is independent of intestinal luminal colonization exclusion and is possible through modulating inflammation.

At the cellular level, the intestinal tract of NE-inflicted birds displays severe small intestinal inflammation, showing massive immune cells infiltration into lamina propria, villus breakdown, and crypt hyperplasia (40, 41). Intestinal inflammation is critical to clear invaded microbes and to resolve inflammation, while overzealous inflammation causes more bacterial invasion and further collateral damage and inflammation (42). Infectious bacteria often hijack the inflammatory pathways to gain survival and invasion advantage. For example, *Salmonella* Typhimurium induces extensive intestinal inflammation and thrives on the inflammation (43). Consistent with this “over-inflammation” model, NE birds with severe intestinal inflammation shows extensive BW loss, villus cell death, and *C. perfringens* invasion. Conversely, DCA attenuating ileitis improves the growth performance and dramatically reduces the NE pathology. Consequently, blocking downstream inflammatory COX signaling by aspirin alleviates intestinal inflammation, villus apoptosis and 60% NE-induced BW loss. These findings indicate that DCA inhibiting inflammatory signaling pathways and targeting them could effectively prevent NE.

Altogether, our data reveal that the microbial metabolic product secondary bile acid DCA attenuates NE, through blunting NE-induced host inflammatory response. These findings highlight the importance of elucidating the molecular relationship between infectious pathogen, microbiome, and host response. These discovers lay the first stone on using microbiome and host inflammatory response to control NE and other intestinal diseases.

## Materials and Methods

### Chicken experiment

All animal protocols were approved by the Institutional Animal Care and Use Committee of the University of Arkansas at Fayetteville. Cohorts of 13 day-of-age broiler chicks obtained from Cobb-Vantress (Siloam Springs, AR) were neck-tagged and randomly placed in floor pens with a controlled age-appropriate environment. The birds were fed a corn-soybean meal-based starter diet during 0-10 days of age and a grower diet during 11-26 days of age. The basal diet was formulated as described before (44). Treatment diets were supplemented with 1.5 g/kg CA or DCA or 0.12 g/kg aspirin (all from Alfa Aesar). Birds were infected with 20,000 sporulated oocytes/bird *E. maxima* at 18 days of age and 10^9^cfu/bird *C. perfringens* at 23 and 24 days of age. Chicken body weight and feed intake were measured at d 0, 18, 23, and 26 days of age. Bird health status was monitored daily after the pathogen infection. Birds were sacrificed at 26 days of age. Ileal tissue and digesta samples were collected for RNA and DNA analysis. Ileal tissue was also Swiss-rolled for histopathology analysis. Images were acquired using a Nikon TS2 fluorescent microscope. Ileal inflammation was scored by evaluating the degree of lamina propria immune cell infiltration, villus shortening, edema, necrosis, crypt hyperplasia, ulceration, and transmural inflammation using a score from 0 to 16.

### *C. perfringens*-induced inflammatory response using primary splenocytes

Splenocytes were isolated similarly to described previously (34). Briefly, chickens at 28 days of age were sacrificed and spleens were resected and homogenized using frosted glass slides in RPMI 1640 medium supplemented with 2% fetal bovine serum, 2mM L-glutamine, 50 μM 2-mercaptoethanol. After lysed the red blood cells, the collected cells were plated at 2 × 10^6^ cells/well in 6-well plates. The cells were pre-treated with 1.2 mM aspirin for 45 min. Cells were then challenged with murine INFγ (1μg/ml, Pepro Tech), TNFa (5 ng/ml, Pepro Tech), or *C. perfringens* (multiplicity of infection 100). The cells were lysed in TRIzol (Invitrogen) at 2 or 4 hours after cytokines or *C. perfringens* treatment, respectively.

### Real time RT-PCR

Total RNA from ileal tissue or splenocytes was extracted using TRIzol as described before (28, 45). cDNA was prepared using M-MLV (NE Biolab). mRNA levels of proinflammatory genes were determined using SYBR Green PCR Master mix (Bio-Rad) on a Bio-Rad 384-well Real-Time PCR System and normalized to *Gapdh*. Ileal digesta DNA was extracted as described before (28) and the digesta bacteria were subject to real-time PCR. The PCR reactions were performed according to the manufacturer’s recommendation. The following gene primers were used: *Cp16S*_forward: 5’- CAACTTGGGTGCTGCATTCC-3’; *Cp16S* reverse: 5’- GCCTCAGCGTCAGTTACAG-3’; *Mmp9*_forward: 5’-CCAAGATGTGCTCACCAAGA-3’ *Mmp9*_reverse: 5’-CCAATGCCCAACTTCTCAAT-3’; *Litaf (Tnfα)*_forward: 5’- AGATGGGAAGGGAATGAACC; *Litaf (Tnfα)*_reverse: 5’-GACGTGTCACGATCATCTGG-3’; *Il1β*_forward: 5’-GCATCAAGGGCTACAAGCTC-3’; *Ill β*_reverse: 5’-CAGGCGGTAGAAGATGAAGC-3’; *Infγ*_forward: 5’- AGCCGCACATCAAACACATA-3’; *Infγ*_reverse: 5’-TCCTTTTGAAACTCGGAGGA-3’; *Ptgs2*_forward: 5’-ACCAGCATTTCAACCTTTGC-3’; *Ptgs2*_reverse: 5’-CCAGGTTGCTGCTCTACTCC-3’; *Gapdh*_forward: 5’-GACGTGCAGCAGGAACACTA-3’; *Gapdh*_reverse: 5’- CTTGGACTTTGCCAGAGAGG-3’.

### TUNNEL assay

Cell apoptosis in intestinal tissue sections was visualized using TUNNEL assay. Briefly, ileal tissue sections were deparaffinized with xylene bath for 3 times and rehydrated with 100%, 95%, and 70% ethanol. The tissue was then incubated with TUNNEL solution (5 μM Fluorescein-12-dUTP (Enzo Life Sciences), 10 μM dATP, 1 mM pH 7.6 Tris-HCl, 0.1 mM EDTA, 1U TdT enzyme (Promega) at 37° C for 90 min. The slides were counter-stained with DAPI for nucleus visualization. The fluorescent green apoptosis cells were evaluated and imaged using a Nikon TS2 fluorescent microscopy.

### Fluorescence in situ hybridization (FISH)

C. *perfringens* at ileal tissue sections was visualized using FISH assay similarly as previously described. Briefly, tissue sections were deparaffinized, hybridized with the FISH probe, washed, stained with DAPI, and imaged using a Nikon TS2 fluorescent Microscope system. The FISH probe sequence of Cp85aa18: 5’-/Cy3/TGGTTGAATGATGATGCC-3’ (46) was used to probe the presence of *C. perfringens* similar to a previous report (45). Briefly, deparaffinized, formalin-fixed 5 μm thick sections were incubated for 15 minutes in lysozyme (300,000 Units/ml lysozyme; Sigma-Aldrich) buffer (25 mM Tris pH 7.5, 10 mM EDTA, 585 mM sucrose, and 0.3 mg/ml sodium taurocholate) at room temperature and hybridized overnight at 46 °C in hybridization chambers with the oligonucleotide probe (final concentration of 5 ng/μl in a solution of 30 percent formamide, 0.9 M sodium chloride, 20 mM Tris pH 7.5, and 0.01% sodium dodecyl sulfate). Tissue sections were washed for 20 minutes at 48 °C in washing buffer (0.9 M NaCl, 20 mM Tris pH 7.2, 0.1% SDS, 20% Formamide, and 10% Dextran Sulfate) and once in distilled water for 10 seconds. The slide was stained with DAPI for 2 min and dried at RT, mounted with 50% glycerol. *C. perfringens* in intestinal tissue was evaluated and imaged using a Nikon TS2 fluorescent microscopy.

### Bile acid C. perfringens inhibition assay

C. *perfringens* in Tryptic Soy Broth (TSB) supplemented with 0.5% sodium thioglycollate, with added TCA, CA or DCA (0, 0.01, 0.05, 0.1, or 0.2 mM, final concentration) or LCA or UDCA (0, 0.2, or 1 mM, final concentration) was cultured overnight under anaerobic conditions. The bacterial growth was monitored by (OD600nm) using a spectrophotometer.

### Statistical Analysis

Values are shown as mean ± standard error of the mean as indicated. Differences between groups were analyzed using the nonparametric Mann–Whitney *U* test performed using Prism 5.0 software. Experiments were considered statistically significant if *P* values were <0.05.

## Acknowledgement

The authors would like to thank the support from staff at Poultry Health Laboratory and Feed Mill in Department of Poultry Science at University of Arkansas, Fayetteville. We also thank P. L. Matsler on helping our histology slides. This research was partially supported by Arkansas Biosciences Institute grant to X. Sun.

